# GenomeDISCO: A concordance score for chromosome conformation capture experiments using random walks on contact map graphs

**DOI:** 10.1101/181842

**Authors:** Oana Ursu, Nathan Boley, Maryna Taranova, Y.X. Rachel Wang, Galip Gurkan Yardimci, William Stafford Noble, Anshul Kundaje

**Affiliations:** Department of Genetics, Stanford University School of Medicine, Stanford, CA, USA, 94305; Department of Statistics, Stanford University, Stanford, CA, USA, 94305; Department of Genome Sciences, University of Washington, WA, USA; Department of Computer Science and Engineering, University of Washington, WA, USA; Department of Computer Science, Stanford University, Stanford, CA, USA, 94305

## Abstract

**Motivation:** The three-dimensional organization of chromatin plays a critical role in gene regulation and disease. High-throughput chromosome conformation capture experiments such as Hi-C are used to obtain genome-wide maps of 3D chromatin contacts. However, robust estimation of data quality and systematic comparison of these contact maps is challenging due to the multi-scale, hierarchical structure of chromatin contacts and the resulting properties of experimental noise in the data. Measuring concordance of contact maps is important for assessing reproducibility of replicate experiments and for modeling variation between different cellular contexts.

**Results:** We introduce a concordance measure called GenomeDISCO (DIfferences between Smoothed COntact maps) for assessing the similarity of a pair of contact maps obtained from chromosome conformation capture experiments. The key idea is to smooth contact maps using random walks on the contact map graph, before estimating concordance. We use simulated datasets to benchmark GenomeDISCO’s sensitivity to different types of noise that affect chromatin contact maps. When applied to a large collection of Hi-C datasets, GenomeDISCO accurately distinguishes biological replicates from samples obtained from different cell types. GenomeDISCO also generalizes to other chromosome conformation capture assays, such as HiChIP.

**Availability:** Software implementing GenomeDISCO is available at https://github.com/kundajelab/genomedisco.

**Supplementary information:** Supplementary data are available at *Bioinformatics* online.

## 1 Introduction

The 3D conformation of chromatin defines a network of physical interactions among genomic loci, including regulatory elements such as gene promoters, distal enhancers and insulators (Krijger and de Laat, 2016). Thus, 3D chromatin architecture plays a key role in gene regulation and cellular function. Changes in 3D chromatin architecture at multiple scales, ranging from large-scale rearrangement of compartments and topologically-associating domains (TADs) to rewiring of enhancer-promoter interactions, are associated with dynamic cellular processes such as differentiation (Fraser *et al.*, 2015; Dixon *et al.*, 2015) and reprogramming (Krijger and de Laat, 2016; Beagan *et al.*, 2016), as well as disease (Lupiáñez *et al.*, 2015; Gröschel *et al.*, 2014).

The last decade has witnessed a revolution in high-throughput sequencing-based assays and imaging techniques to map 3D chromatin architecture at multiple scales and resolutions, providing new insights into spatial genome organization (Schmitt *et al.*, 2016). The sequencing-based methods (referred to as 3C-seq experiments) for assaying 3D chromatin architecture such as 3C (Dekker *et al.*, 2002), 4C (Zhao *et al.*, 2006; Simonis *et al.*, 2006), 5C (Dostie *et al.*, 2006), Hi-C (Lieberman-Aiden *et al.*, 2009), Capture Hi-C (CHi-C) (Mifsud *et al.*, 2015), ChIA-PET (Fullwood *et al.*, 2009) and HiChIP (Mumbach *et al.*, 2016) are all variations of the chromosome conformation capture technique. In a Hi-C experiment, genome-wide interactions are mapped by ligating proximal fragments followed by deep sequencing. The result of such an experiment is a genome-wide contact map, which is a matrix with a sequencing readout of the contact frequency for every pair of genomic loci.

A number of computational methods have been designed to normalize (Yaffe and Tanay, 2011; Hu *et al.*, 2012; Imakaev *et al.*, 2012; Knight and Ruiz, 2013; Servant *et al.*, 2015) and extract statistically significant contacts from the different types of 3D chromatin conformation assays (Ay *et al.*, 2014; Ron *et al.*, 2017; Mifsud *et al.*, 2017; Cairns *et al.*, 2016; Carty *et al.*, 2017). However, principled methods for systematic comparisons of 3D contact maps are equally important and form a core component of two key analyses. First, as an essential quality control tool, it is useful to quantify the concordance of replicate experiments. This is particularly relevant because it is common practice to pool reads across biological replicates of a 3C-seq experiment before downstream analyses. Significant differences between the pooled replicates could result in suboptimal or misleading downstream results. Second, understanding and quantifying similarity between replicates is also an essential step in differential analysis, where the goal is to reliably identify statistically significant differences between contact maps in different biological conditions. Differences between conditions can only be trusted if they exceed the differences between biological replicates.

Experimentally derived contact maps exhibit certain properties that are distinct from other types of functional genomic data. First, contact maps explicitly encode the adjacency matrix of a multi-scale, modular network consisting of large-scale compartments, TADs, CTCF/cohesin mediated loops and potentially transient interactions between other types of elements (Schmitt *et al.*, 2016). Second, the contact frequency between a pair of loci is strongly dependent on their linear genomic distance (Dekker *et al.*, 2002; Ay *et al.*, 2014; Duan *et al.*, 2010) and affected by additional biases such as restriction fragment size, GC content and mappability (Yaffe and Tanay, 2011; Imakaev *et al.*, 2012; Cournac *et al.*, 2012; Hu *et al.*, 2012; Schmitt *et al.*, 2016). Third, the resolution of a contact map defined in terms of the size (in nucleotides) of the interacting loci is often a free parameter and heuristically determined based on the depth of sequencing (Rao *et al.*, 2014). Finally, the noise associated with estimates of contact frequencies is also strongly associated with sequencing depth. These properties necessitate the development of new computational methods specifically suited for analysis of Hi-C data.

Statistical measures that have been developed to quantify the reproducibility of 1D functional genomics assays, such as ChIP-seq, DNA methylation and RNA sequencing, cannot be trivially applied to 3D contact maps. For instance, simple correlation measures, which are most frequently used as measures of reproducibility (Rao *et al.*, 2014), do not correctly capture the reproducibility of Hi-C data (Yang *et al.*, 2017; Yardımcı *et al.*, 2017). This is partly because these simple correlation measures consider each entry in a contact map as an independent measurement, thereby ignoring the rich connectivity and dependence structure in 3D contact maps. More sophisticated reproducibility measures have recently been introduced including comparison of eigenvectors (Yan *et al.*, 2017) and a stratified correlation coefficient (Yang *et al.*, 2017) and these methods alleviate many of the problems with traditional correlation.

In this work, we introduce GenomeDISCO (DIfferences between Smoothed COntact maps), a computational framework for quantifying reproducibility or concordance of contact maps from 3C-seq experiments. We represent a contact map as a network or graph, where nodes are genomic loci and edges are weighted proportional to appropriately normalized contact frequency between a pair of loci (nodes). We denoise the contact maps using random walks on the graph, followed by comparison of the resulting smoothed contact maps. We use systematic simulations to calibrate the method, showing its ability to detect artificially introduced noise, differences in distance dependence curves and differences in structural properties of contact maps. We then apply GenomeDISCO and other related approaches to the largest existing collection of Hi-C experiments (Rao *et al.*, 2014) and benchmark their performance on a comparison of replicate experiments and experiments from different cell types. We also show that GenomeDISCO easily generalizes to other types of 3C-seq assays, such as HiChIP. We provide an efficient implementation of our method as well as comprehensive analysis reports and visualizations in a user-friendly software package at https://github.com/kundajelab/genomedisco. GenomeDISCO is also included in the 3D genome analysis suite recommended by the Encyclopedia of DNA Elements (ENCODE) Consortium at https://github.com/kundajelab/3DChromatin_ReplicateQC (Yardımcı *et al.*, 2017).

## 2 Methods

### 2.1 A graph representation of chromatin contact maps

We represent a contact map as a graph or network of interactions between genomic loci, with adjacency matrix *A*. Each node *i* in the network is a genomic locus (segment) of a specified resolution or size (in nucleotides). The weight of each edge *A*_*ij*_ is a normalized, experimentally-derived contact frequency between a pair of nodes *i* and *j*. In this work, we normalize the contact map using the sqrtvc normalization (for additional discussion of normalization methods compatible with GenomeDISCO, refer to the Supplementary Methods), and convert it to a transition probability matrix, such that all rows sum to 1. This transition matrix is the weighted adjacency matrix A used in the analyses in this study. We ignore inter-chromosomal interactions and hence represent all chromosomes as independent graphs.

### 2.2 Motivation for our concordance score

A concordance score that aims to estimate the global similarity between a pair of contact maps must account for the specific properties of experimentally-derived contact maps. First, contact maps contain structural features that manifest at different scales, such as large scale compartments, sub-Mb scale TADs and sub-TADs that manifest as densely connected diagonal blocks and CTCF/cohesin mediated loops observed as focal points of enriched contacts. Thus, an ideal concordance score would be able to measure similarity across multiple scales. Second, genome-wide contact maps such as those from Hi-C experiments measure a very large space of possible contacts and hence require deep sequencing (> billion reads) for reliable estimates of contact frequency.

**Fig. 1.**
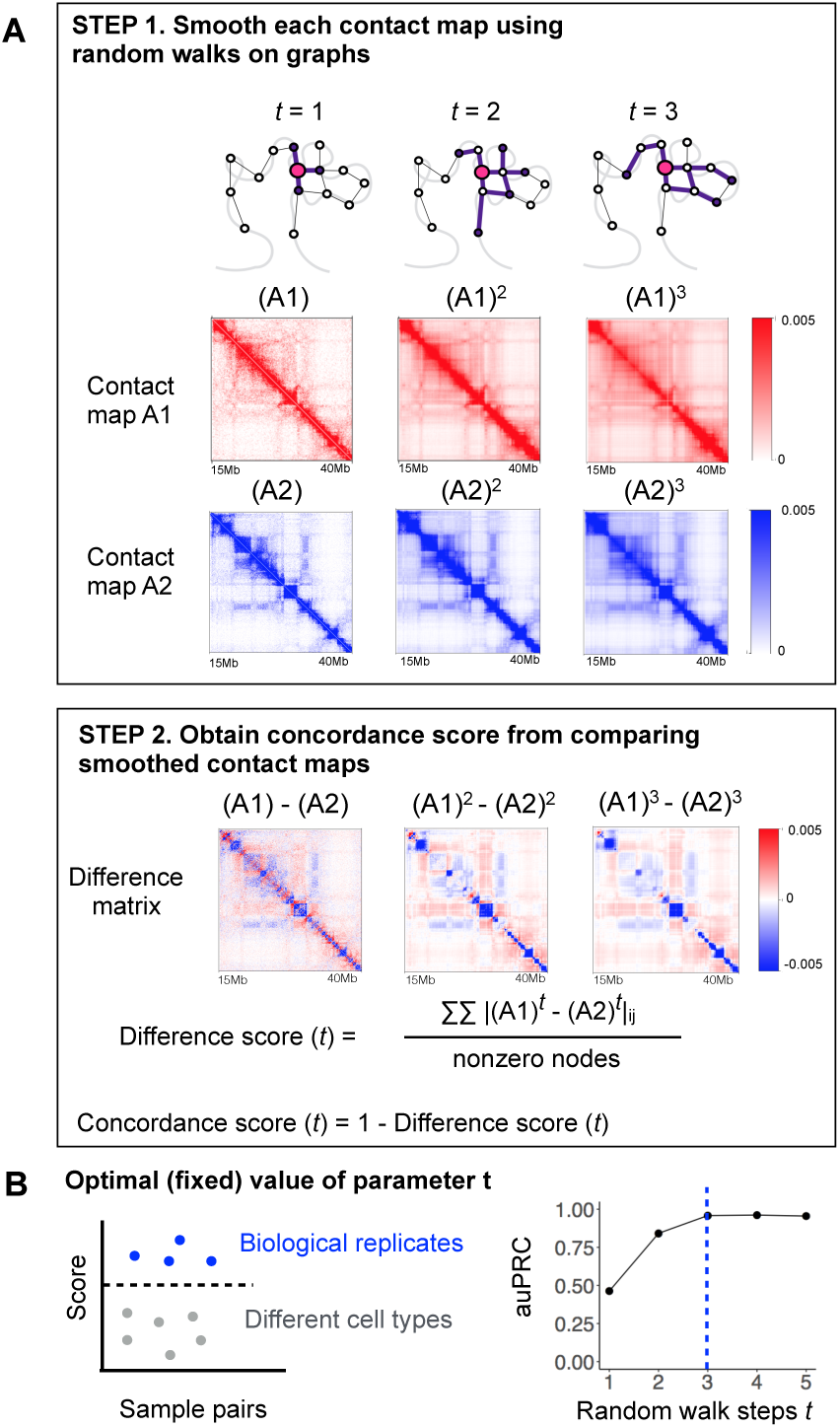
Overview of GenomeDISCO. A) GenomeDISCO consists of 2 steps. The first step in comparing 2 contact maps, *A*1 and *A2*, consists of smoothing each contact map using random walks. Depicted are the smoothed contact maps, at different levels of smoothing controlled by the parameter *t*, which specifies the number of steps of random walk used for denoising. The second step consists of computing a difference score between the smoothed contact maps, as a function of *t* C) Procedure for identifying the optimal value for *t*. We computed concordance scores for pairs of samples that are either biological replicates from the same cell type, or pairs of samples from different cell types. We assume that the optimal value of *t* will produce scores that can accurately classify pairs of samples into “Biological replicates” and “Different cell types”. For each value of *t*, we measure classification performance using the area under the precision-recall curve (auPRC), finding *t* =3 to be optimal.

Due to cost and material constraints, typical Hi-C datasets are sequenced at significantly lower coverage (e.g. 100M reads (Lajoie *et al.*, 2015)). These undersampled datasets exhibit a large proportion of contacts with low observed counts with high variance (Carty *et al.*, 2017) including some contacts with 0 observed counts, a phenomenon known as stochastic dropout. To address this issue, we propose a denoising approach to smooth contact maps by leveraging random walks on the contact map graph, before comparing these maps.

### 2.3 The GenomeDISCO score for estimating the concordance of contact maps

We estimate concordance between a pair of chromatin contact maps, *A*1 and *A2*, as follows (Figure 1A).

#### Equalizing sequencing depth

To avoid artificial differences due to sequencing depth (see Supplementary Figure 2), we first equalize the sequencing depth of the pair of datasets to be compared by randomly subsampling the count matrix to the minimum depth of the two datasets.

#### Denoising contact maps using random walks

We denoise each contact map independently using random walks on the contact maps. For every pair of nodes *i* and *j* in a contact map, we ask the question: if we start a random walk at node *i* based on the observed contact map transition probability matrix, and allow the walk to take *t* steps, what is the probability we will reach node *j*? If there are many high-probability paths in the network that connect node *i* and node *j*, it increases our confidence that node *i* and node *j* are in contact. The probability of reaching node *j* after a random walk of *t* steps starting from node *i* is the (*i,j*)th entry of the matrix obtained by multiplying the transition probability matrix with itself *t* times i.e. (*A*^*t*^)_*i,j*_. We define the optimal value for the steps parameter *t* for Hi-C data, as the one that maximizes the ability of the concordance score to distinguish between biological replicates and non-replicate reference datasets (See Section 2.4 for details).

#### Computing the difference between denoised contact maps

The denoised versions of contact maps *A*1 and *A*2, after *t* steps of random walk are (*A*1^*t*^) and (*A*2^*t*^) respectively. We compute the difference *d*_*t*_(*A*1,*A*2) between *A*1 and *A*2 as the L_1_ distance between the two denoised contact maps (*A*1^*t*^) and (*A*2^*t*^), divided by the average number of non-zero nodes in the two original contact maps *A*1 and *A*2:

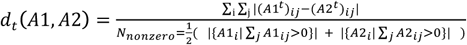

Since each row of *A*1 and *A*2 sums to 1, the weighted degree (sum of weights of all edges to/from a node) of each node is 1. Hence, *d*_*t*_(*A*1, *A*2) scores range from 0 to 2, with small values indicating high similarity.

#### Converting the difference to a concordance score

We define the concordance score as *R*(*A*1, *A*2, *t*) *= 1- d*_*t*_(*A*1, *A*2). The concordance scores range from −1 to 1, with larger values indicating greater similarity. We obtain a single genome-wide score as the average of the scores across all chromosomes.

### 2.4 Estimating the optimal number of random walk steps (*t*)

The number of steps *t* of the random walk on the contact map graph determines the amount of smoothing or denoising of a contact map. We define an optimal value of *t* as one that would provide sufficient denoising so as to improve concordance between contact maps of replicate experiments while preserving differences between contact maps from distinct cellular contexts. We used a collection of high quality benchmark Hi-C datasets with replicate experiments from diverse human celllines (Rao *et al.*, 2014) to optimize *t*. Using half the experiments as a training set and the remaining half as a test set, we asked which value of *t* leads to the optimal separation of biological replicates from non-replicate samples, as measured with the area under the precision-recall curve (auPRC). We found *t* = 3 achieved the best performance on the training set (auPRC of 0.95, Figure 1B), associated with an auPRC of 0.92 on the test set. The optimal value of *t* = 3 identified using reference Hi-C datasets generalized to HiChIP data (see Figure 4) and to Hi-C datasets from other species such as *Drosophila* (see (Yardımcı *et al.*, 2017)). It is possible that for other applications of GenomeDISCO, other values of *t* may be optimal. In such cases, we suggest users perform a similar calibration experiment to identify the optimal value.

**Fig. 2.**
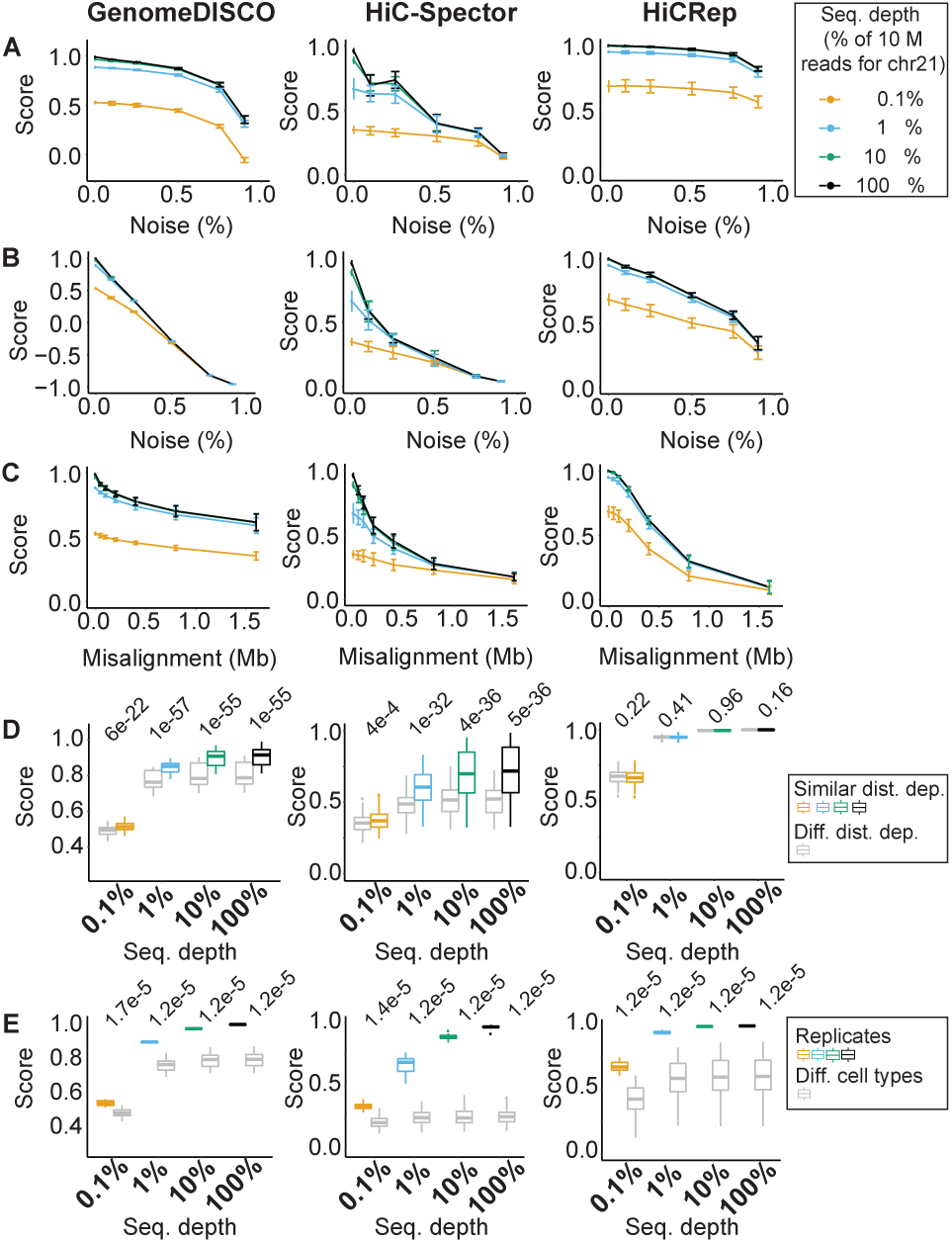
GenomeDISCO exhibits desired features for a reproducibility score. A-D) Scores as a function of edge dropout (A), node dropout (B), domain boundary misalignment (C) and difference in distance dependence curves (D) for GenomeDISCO, HiC-Spector and HiCRep methods. For A-D) error bars represent one standard deviation from the mean score, based on independent simulations across all cell types from (Rao *et al.*, 2014). For D) we split pairs of samples into “similar distance dependence” and “different distance dependence” using a threshold of 0.005 Jensen Shannon divergence between the curves of the samples compared (see Supplementary Methods). E) Results on simulations comparing replicates with non-replicates obtained from different cell types. For D-E), values above the plots are p-values of a Mann-Whitney test.

## 3 Results

### 3.1 Benchmarking GenomeDISCO on simulated perturbations to 3C-seq datasets

We expect an effective concordance score for 3C-seq datasets to be sensitive to key types of noise and artifacts that typically affect these data (Supplementary Figure 1).

We benchmarked the behavior of GenomeDISCO by using it to compute concordance between a reference Hi-C contact map and a version of the map that is explicitly perturbed with different types and levels of simulated noise (See Supplementary Methods). We performed our analyses at 50kb resolution, as this is a resolution frequently used in the analysis of Hi-C datasets. We compared GenomeDISCO to two other recently developed methods for estimating concordance of Hi-C data: HiCRep, which measures correlation of contacts stratified by distance (Yang *et al.*, 2017), and HiC-Spector, which computes an eigendecomposition of the Laplacian of the graph, and then compares the L_2_ distance between eigenvectors of the 2 contact maps (Yan *et al.*, 2017).

We examined the sensitivity of the concordance scores to perturbations that involve random dropout of edges and nodes as well as misalignment of domain boundaries in the perturbed contact map relative to the reference. Indeed, we found that concordance scores from all three methods decrease with increasing edge drop out (Figure 2A), increasing node drop out (Figure 2B) and increasing domain boundary misalignment (Figure 2C, see Supplementary Methods).

Next, in order to understand the effect of sequencing depth of the contact maps, we repeated the above three perturbation analyses for reference and perturbed maps subsampled to four depths: 100%, 10%, 1%, 0.1% of 10 million reads restricted to chromosome 21. As expected, we found that the GenomeDISCO score was the highest for the most deeply sequenced samples. Concordance scores dropped consistently with decreasing sequencing depth across all types and levels of perturbations (Figure 2). The scores were found to plateau as the sequencing depth increased from 1 million to 10 million reads, which is expected, since for a 50kb resolution, one would need approx. 0.8 million reads for chr21 (see Supplementary Methods).

Contact maps can also differ in their fundamental distance dependence curves that capture the probability of contact as a function of linear genomic distance. Distance dependence curves have been found to change due to cell cycle stage (Naumova *et al.*, 2013; Nagano *et al.*, 2017) or as a function of perturbation of proteins involved in chromatin 3D architecture, such as RAD21 knockout in yeast (Mizuguchi *et al.*, 2014) or WAPL and SCC4 knockouts in human HAP1 cells (Haarhuis *et al.*, 2017). Replicates from the same condition are often pooled, and if they have different distance dependence curves, the result will be an average that is not representative of either replicate. Hence, being sensitive to differences in distance dependence curves is a useful property of a concordance score.

We simulated pairs of contact maps from a common reference contact map by sampling reads according to two different distance dependence curves, obtained from Hi-C maps from pairs of different cell types (see Supplemental Methods). We split the pairs of contact maps into pairs with similar distance dependence curves and pairs with different curves (see Supplementary Methods), and compared the scores we obtained at different sequencing depths (as above) using all three methods. GenomeDISCO samples with different distance dependence curves obtain lower concordance scores. As in the other simulations, the margin between the two sets of pairs decreased as we decreased sequencing depth (Figure 2D). HiC-Spector was also sensitive to differences in distance dependence curves, while HiCRep was not. GenomeDISCO had the best margins of separation at lower sequencing depths.

Finally, we asked whether pairs of simulated pseudo-replicates sampled from the same reference Hi-C map would be deemed more concordant than pairs of samples from different cell types. All three methods successfully discriminated the two sets of pairs with margins decreasing with decreasing sequencing depth (Figure 2E).

### 3.2 Benchmarking GenomeDISCO on Hi-C datasets

We used more than 80 high quality Hi-C datasets from (Rao *et al.*, 2014) spanning multiple human cell-lines (GM12878, HMEC, HUVEC, IMR90, K562, KBM7, NHEK) to benchmark the behavior of our concordance score (Figure 3, Supplementary Table 1, 2). Due to the lack of explicit ground truth about the nature of noise in real datasets, we evaluate the validity of the concordance score by expecting higher scores when comparing pairs of biological replicates of Hi-C data with similar distance-dependence characteristics as compared to scores obtained by comparing Hi-C datasets from different cell types. We focused our analysis on a subset of experiments defined as those done with in-situ Hi-C (see Supplementary Table 2).

**Fig. 3.**
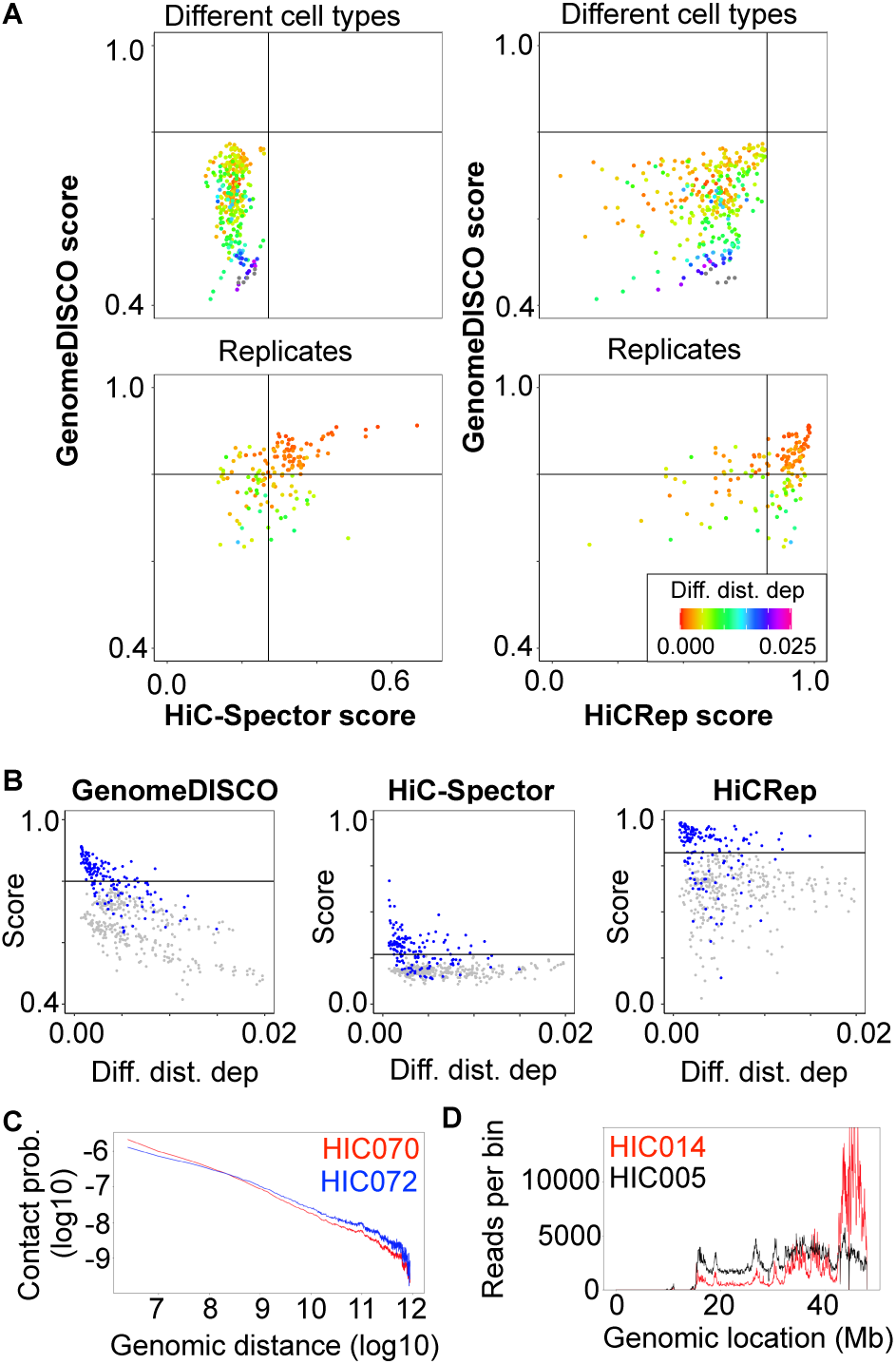
GenomeDISCO distinguishes biological replicates from nonreplicates, taking distance dependence curves into account. A) Scatterplot of scores obtained with GenomeDISCO vs those obtained with HiCRep and HiC-Spector. For each of the three methods, we define a threshold that separates low-concordance from high-concordance pairs of samples. The threshold is chosen as the highest score obtained by a comparison between different cell types. GenomeDISCO largely agrees with the other methods. There is a subset of scores that GenomeDISCO selectively ranks as low-concordance, and those involve pairs of contact maps with large differences between their distance dependence curves. B) Concordance scores as a function of difference in distance dependence functions. The difference is measured as the Jensen Shannon divergence between the contact probability distributions (see Supplementary Methods). C) Example of different distance dependence functions that GenomeDISCO deems non-concordant but HiCRep defines as concordant. D) Row sums for each genomic bin for sample HIC014 are non-uniform, compared to e.g. HIC005, at a similar sequencing depth of ∼ 300 million reads.

Next, we used GenomeDISCO, HiCRep and HiC-Spector to compute concordance scores for all the pairs of biological replicates and pairs of samples from different cell types. Hierarchical clustering of the samples based on the matrix of all pairwise concordance scores revealed that samples from the same cell type cluster together, for all three methods (see Supplementary Figure 5). For each method we defined an empirical threshold for classifying sample-pairs into one of two categories labeled high-concordance and low-concordance. The threshold was determined as the highest score across all pairs of samples from different cell types, since we expect concordant biological replicates to be at least as concordant as samples from different cell types. We then analyzed the similarities and differences between the three methods in terms of their classification of the pairs of biological replicates. (Figure 3A).

Out of 149 pairs of biological replicates in the test set, we found that the methods agreed across most samples (94/149 biological replicate pairs were classified consistently between GenomeDISCO and HiCRep, and 102/149 between GenomeDISCO and HiC-Spector). For a small subset of replicate-pairs, HiCRep and/or HiC-Spector classified them as high-concordance, while GenomeDISCO classified them as low concordance: of these 21/34 of the comparisons deemed concordant by HiCRep and 12/23 by HiC-Spector, the comparisons involved samples with large differences in distance dependence curves (difference in distance dependence curve higher than 0.005, a value that was found to distinguish pairs of biological replicates in the high-concordance class from those in the low concordance class). For example, samples HIC070 and HIC072 (biological replicates for the K562 cell line) are classified as low-concordance by GenomeDISCO (score 0.644), but classified as high-concordance by HiCRep (score 0.910). These samples have a marked difference in their distance dependence curves (ranked as the largest difference in distance dependence curve among all biological replicate pairs) (Figure 3C). In fact, GenomeDISCO scores in general drop proportional to the difference in distance dependence curves between the pair of samples being compared (Figure 3B). Finally, we find 18 cases ranked as non-concordant by both HiCRep and HiC-Spector but deemed concordant by GenomeDISCO. For 6/18 of these, the GenomeDISCO score is equal to the threshold concordance of 0.8. Similarly, there are 18 cases deemed concordant only by HiCRep and 7 deemed concordant only by HiC-Spector.

We also found that 18 replicate pairs were deemed low-concordance by all three methods. In particular, in eight of these cases, replicate pairs classified as low-concordance by all three methods involved sample HIC014 from the GM12878 cell type (specifically HIC014 vs any of HIC004, HIC006, HIC010, HIC018, HIC022, HIC038, HIC042, HIC048). Upon closer inspection, we found that HIC014 exhibited an unusual pattern of uneven coverage across the genome (Figure 3D), likely explaining the observed results.

Finally, we also used the Hi-C data to check whether GenomeDISCO is able to detect differences in protocols or restriction enzymes used for each experiment (see Supplementary Figure 4A). We found that GenomeDISCO scores are lower for comparisons between samples prepared with dilution Hi-C versus in situ Hi-C. This observation is expected because dilution Hi-C experiments capture more random ligations between nuclear and mitochondrial DNA than in-situ Hi-C (see (Rao *et al.*, 2014)). We also found that GenomeDISCO scores are higher for experiments performed with the same enzyme, compared to different enzymes (Supplementary Figure 4B).

### 3.3 Benchmarking GenomeDISCO on HiChIP data

We applied GenomeDISCO to a set of H3K27ac HiChIP datasets from (Mumbach *et al.*, 2017), covering 2-3 replicates for 7 cell types (GM12878, HCASMC, K562, My-La and 3 types of T-cells: Naïve, Th17 and Treg, see Supplementary Table 1). As for Hi-C, we binned the HiChIP reads at a resolution of 50 kb and normalized the contact maps using sqrtvc. We then ran GenomeDISCO on all pairwise comparisons and checked whether biological replicates are deemed more concordant than pairs of samples from different cell types. We found that GenomeDISCO scores correctly separate biological replicates from nonreplicates, for the non-T-cell comparisons, using the same parameters as for Hi-C and the same threshold for defining concordance (from Figure 3), suggesting that GenomeDISCO generalizes seamlessly to HiChIP data (Figure 4). We obtained similar results for HiCRep and HiC-Spector. For the comparisons between T-cells, all three methods produced similar scores for both comparisons between biological replicates and those between different types of T-cells, with the biological replicates receiving the highest scores in almost all cases. Using the thresholds of concordance derived for Hi-C, we find that for GenomeDISCO, T-cell related comparisons pass the threshold above a sequencing depth of 50 million reads, while HiCRep deems all T-cell comparisons as concordant and HiC-Spector deems a smaller subset as concordant. Overall, we find that GenomeDISCO behaves as expected for HiChIP data, without any modifications to the method.

**Fig. 4.**
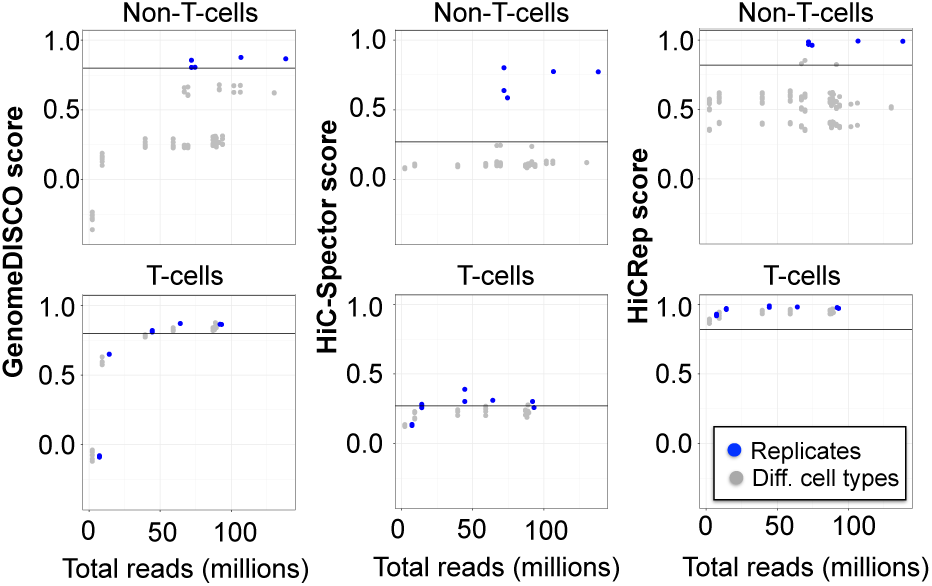
GenomeDISCO benchmarks on HiChIP data. A) Scores obtained on HiChIP data for GenomeDISCO, HiC-Spector and HiCRep. The scores are split into two categories: comparisons between T-cells and the remaining comparisons (labeled as “Non T-cells”). Scores of replicates (in blue) are plotted offset by 5 million reads, to improve visibility of points that would otherwise overlap.

## 4 Discussion

Here, we present GenomeDISCO, a new approach specifically designed for evaluating concordance and reproducibility of chromatin contact maps. Our benchmarking experiments on simulated contact maps and high quality real Hi-C and HiChIP datasets, which include systematic comparisons to two other methods HiCRep and HiC-Spector, indicate that GenomeDISCO displays competitive accuracy in distinguishing biological replicates from different cell types with the desired sensitivity to sequencing depth, node and edge dropout noise, changes in domain boundaries and subtle differences in distance dependence.

GenomeDISCO introduces a novel approach of using random walks on the contact map graph for progressive smoothing and evaluation of concordance at multiple scales. A weighted graph is a natural representation of a chromatin contact map. A random walk on a contact map graph progressively upweights direct edges involving node pairs that have many high-weight indirect paths of progressively increasing lengths that connect the node pairs.

Further, GenomeDISCO is sensitive to subtle differences in distance dependence curves. Since it is common to pool multiple Hi-C replicates, it is essential to know if samples exhibit differences, so as to not eliminate signal during pooling, especially since in some cases variation in distance dependence curves is biologically meaningful.

On the other hand, two datasets can have different distance dependence curves but still be concordant in terms of enrichments of contacts when accounting for the different distance dependence function of each dataset. Thus, if one is interested in evaluating concordance of contact enrichment (e.g. as measured by methods that call significant contacts), then one can normalize the observed contact frequencies by the expected distance-dependent contact frequencies (which would correct for most differences in distance dependence) for the pair of contact maps before feeding them into GenomeDISCO. One can obtain these observed/expected ratios or associated *q*-values from Fit-Hi-C (Ay *et al.*, 2014).

Further, GenomeDISCO provides a variety of diagnostic analyses that are useful for digging deeper in the potential reasons for low concordance. The diagnostic analyses include the comparison of distance dependence curves, and a difference matrix between smoothed contact maps (Figures 1 and 3).

Finally, what determines a good threshold for concordance of biological replicates? Based on our extensive analyses of simulated datasets and extensive collections of Hi-C data, we define an empirical GenomeDISCO score threshold of 0.8 at 50kb resolution. We also provide a set of precomputed standards based on pseudo-replicates for frequently used resolutions, allowing a direct calibration of a given score to an upper bound.

While GenomeDISCO summarizes concordance in a single score, a future direction of research consists of developing methods that specifically focus on measuring concordance of distinct features of the contact map, such as TADs, compartments and loops. For cases where concordance is low, such methods will be instrumental to pinpoint the specific feature of the contact maps that accounts for the observed difference. Three-dimensional chromatin architecture is the next frontier in deciphering genome function. Ensuring high quality reproducible experiments is an essential part of this revolution in understanding chromatin architecture. GenomeDISCO is a user-friendly, efficient and accurate diagnostic tool for evaluating the reproducibility of 3D chromatin conformation capture experiments.

## Author contributions

A. K., O. U conceived GenomeDISCO. O. U. implemented GenomeDISCO, designed benchmarks, and ran experiments. N. B, M. T., R. W., G. Y. and W. S. N. provided useful discussions. O. U., A. K. wrote the paper. O. U., A. K., N. B, M. T., R. W., G. Y. and W. S. N edited the paper.

## Acknowledgements

We thank Michael Snyder, Jonathan Pritchard, Howard Chang, Michael Bassik, Avanti Shrikumar and the Kundaje Lab for helpful discussions. We thank Suhas Rao for clarifications related to the Hi-C datasets used and Anna Shcherbina and Chris Probert for help with visualization.

## 1 Datasets, preprocessing and normalization

### Hi-C Data

We used Hi-C datasets from seven cell types from (Rao *et al.*, 2014) as summarized in Supplementary Table 1 (GEO accession numbers included in the table). For each cell type, we downloaded reads mapped to the hg19 human genome reference and filtered them for mapping quality (MAPQ > 30). We then computed the number of reads supporting contacts between all pairs of genomic bins of a specified resolution to obtain a contact map. In general, GenomeDISCO expects a userdefined resolution, which can be determined empirically, for instance using the definition provided in (Rao *et al.*, 2014), i.e. choosing the lowest resolution such that at least 80% of the genomic bins have at least 1000 contacts with non-zero counts. In this work, we used a 50 kb resolution for all analyses, as this resolution is typically used in the analysis of Hi-C datasets.

### HiChIP Data

We used HiChIP datasets from seven cell types from (Mumbach *et al.*, 2017), as summarized in Supplementary Table 1 (which includes GEO accession numbers). We downloaded “allValidPairs” files from GEO, and binned them at 50 kb resolution to obtain raw contact maps.

### Normalization of Hi-C and HiChIP data

For both Hi-C and HiChIP data, we started from a raw contact map *C* and obtained its normalized version *A* as follows.

First, we normalized *C* using the square root normalization method (sqrtvc from (Rao *et al.*, 2014)) that corrects for node-specific, factorizable biases:

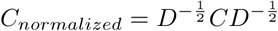

where *D* is a diagonal matrix, with each entry *D*_*ii*_ corresponding to the degree (row sum) of node *n*_*i*_. Other normalization frameworks such as ICE (Imakaev *et al.*, 2012) or KR (Knight and Ruiz, 2013) are also compatible with our framework and do not change any presented conclusions. We prefer the sqrtvc normalization since KR and ICE occasionally do not converge for Hi-C datasets with moderate sequencing depth processed at very high resolution, such as 5-10kb.

Second, to obtain *A* from *C*_*normalized*_, we rescale the values in each row of *A*, such that all rows of *A* sum to 1 (i.e. *A* is a valid transition matrix).

The final transition matrix *A* is the input to the random walk denoising described in the main text.

**Note about additional normalization due to copy number variation**

The goal of GenomeDISCO is to estimate the concordance between two samples, processed as they will be used for downstream analysis. This is why we normalize datasets before performing the smoothing and the subsequent comparison. If for downstream analyses, a different normalization scheme is necessary (such as correcting for copy number variation), we suggest that users perform their desired normalization and then run GenomeDISCO on the normalized data. In such a case, the sqrtvc normalization that GenomeDISCO does by default should be turned off.

## 2 GenomeDISCO

### 2.1 A network representation of contact maps

We consider the contact map as a network, where nodes correspond to genomic bins, and edges are weighted by the contact frequency in the contact map, as described in the main text section 2.1.

### 2.2 Desired features of a concordance score for 3C-seq data

We reasoned that an effective concordance score must account for multiple properties of 3C-seq data, as illustrated in Figure S1.

First, the score should decrease if we randomly drop out edges of the contact map. This is illustrated in Figure S1A, where we compare a contact map (in red) with the same contact map that has a fraction of its edges randomly removed. Similarly, we expect that random removal of nodes would also reduce concordance (illustrated in Figure S1B), as would differences in the positions of the domains in the contact maps (illustrated in Figure S1C). In addition, we expect a sound concordance score to rank replicate experiments as more concordant than pairs of experiments performed in different cell types or different conditions (Figure S1D).The score should also measure differences in the distance dependence of contact probability (Figure S1E). Finally, we expect better concordance at higher sequencing depth for replicate experiments (Figure S1F), as more features of the contact map should be recovered in each experiment.

### 2.3 Motivation for subsampling datasets to equal sequencing depth before computing concordance scores

The first step when computing GenomeDISCO scores consists of subsampling the pair of datasets to the sequencing depth of the dataset with lower coverage. This is done for two reasons.

First, we bring datasets to equal sequencing depths because we want to be conservative in the concordance scores we provide. We assume that the maximum concordance of the pair of data is based on the lowest sequencing depth of the 2 datasets. In other words, a dataset sequenced at low sequencing depth cannot capture the rich information of a dataset with higher coverage.

Second, we downsample datasets in order to make it possible for users to make comparisons *between* concordance scores. Since GenomeDISCO scores generally increase with additional sequencing, they cannot be directly compared without accounting for sequencing depth. To illustrate this, we have performed the following experiment: we took the existing datasets from 7 cell types profiled in Rao *et al.* (2014), and performed all pairwise comparisons between these datasets, but with each dataset subsampled to either 100000 or 10000 reads for chr21. In this way, we can compare the scores we obtain when each dataset is considered at either a high (100000) or low (10000) sequencing depth, and inspect what happens when we do not perform subsampling to the same depth. The results are presented in Figure S2.

Consider for example, the pair of samples GM.10000.rep1 and GM.1000.rep2, which are simulated replicates from the GM12878 cell type, both with a low sequencing depth. When they are compared to each other, they get a score of 0.4. However, comparing either one of the GM.10000 replicates to a more deeply sequenced sample from a different cell line, e.g. K562.100000.rep2 gives a score of 0.5. The higher score of 0.5 obtained for the cross-cell type comparison compared to the 0.4 score obtained for replicates may give the wrong impression that the samples from different cell types are more similar than those from the same cell type. However, the comparison against K562 at the same sequencing depth gives the score 0.35, which is as expected lower than that of biological replicates measured at that sequencing depth.

It is important to note that, although we subsample datasets for the purposes of computing concordance scores, we are not suggesting that datasets cannot be used for downstream analyses at their full sequencing depths. For instance, analyses that do not involve comparing samples across conditions (such as calling domains, compartments, loops in a single dataset) certainly benefit from using datasets at their highest coverage. We are only suggesting the subsampling for cross-condition comparisons, where variation in sequencing depth may lead to detection of artificial differences due to the dataset with lower coverage missing features present in the dataset with higher coverage.

**Figure S1:**
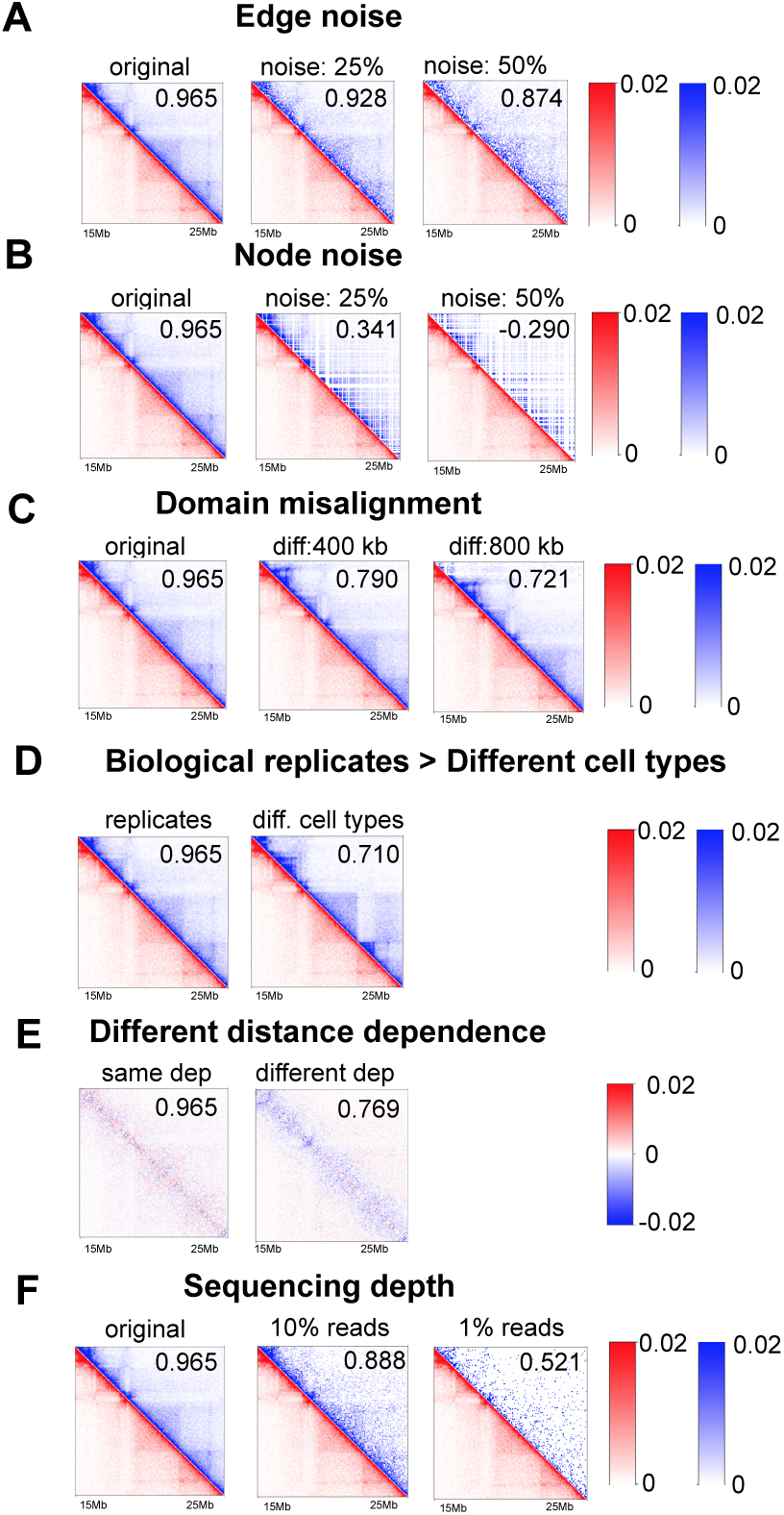
Desired features of a concordance score for 3C-seq data. A good concordance score should be sensitive to edge dropout (A), node dropout (B), misalignment of domain boundaries (C), differences in distance dependence curves (D) and differences in contact frequency such as those observed between different cell types (E). It should also decrease as sequencing depth is reduced (F). For A),B),C),D) and F), we plot pairs of contact maps, where in red we show the original contact map, and in blue we show the contact map perturbed as described above. For E), we plot the difference matrix obtained between datasets with either the same or different distance dependence curves. For A-E), we include the score computed by GenomeDISCO for each pair of contact maps in the upper right corner of the plotted matrices.

**Figure S2:**
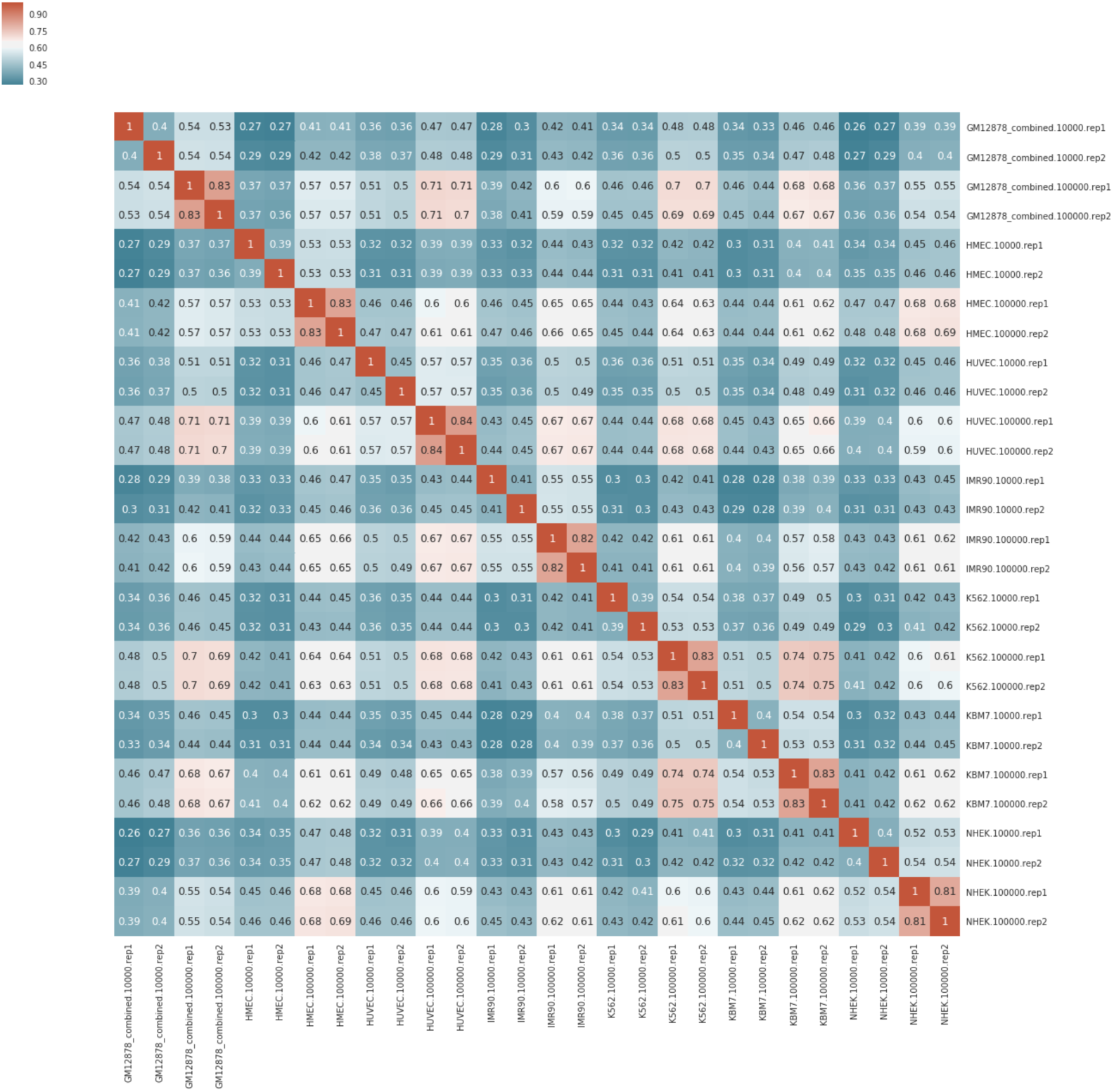
Subsampling experiment.

### 2.4 Calibrating the concordance score

We calibrated the concordance scores using biological replicates as the gold standard for defining similarity and datasets from different cell types as the gold standard for defining dissimilarity. Specifically, for Hi-C data, we identified concordance score thresholds that best distinguished pairs of simulated replicates from pairs of simulated datasets representing contact maps from different cell types (See Section 3.1). Concordance scores also depend on the baseline resolution of the contact maps (higher resolutions result in an overall decrease in the magnitude of concordance scores). Hence, we provide precomputed calibrated thresholds for concordance scores for a range of resolutions: 10 kb, 50 kb, 500 kb at https://github.com/kundajelab/genomedisco under the directory “calibration_tables”.

## 3 Benchmarking GenomeDISCO on simulated data

### 3.1 Simulating different types of noise in contact maps

We first simulated realistic contact maps based on Hi-C datasets from seven cell types from (Rao *et al.*, 2014) (see Supplementary Table 1 for a list of datasets used for the simulations) in order to calibrate parameters and evaluate the performance of GenomeDISCO. We pooled reads across all replicates for each cell type. We used a resolution of 50 kb fixed size genomic bins. For the sake of efficiency, we then restricted the contact map to chromosome 21 for our simulations. For each cell type, we rescaled the raw contact count matrix *C* into a probability matrix *P*, such that all entries in the upper triangle of *P* sum to 1 (i.e. a valid probability distribution). Then, to distinguish structural differences (domains, compartments) from differences in distance dependence curves, we scaled the obtained probabilities to ensure that datasets for all cell types contain identical distance dependence functions. We modified the distance dependence curves to follow one reference distance dependence function. For the Hi-C datasets, we used the GM12878 dataset as the reference since it is the most deeply sequenced cell type. For each genomic distance *g* (from 0 bp to the length of the chromosome), we divided each entry at genomic distance *g* by the ratio of the average contact probability at distance *g* in the target dataset and the average contact probability at distance *g* in the reference dataset with the desired distance dependence curve. Note that the upper triangle of the resulting scaled matrix is a valid probability distribution because the upper triangle of the reference matrix is a probability distribution. Finally, we simulated a contact map of a desired read depth *N* by sampling each entry (*i, j*) from a binomial distribution, with *p* = *P*_*ij*_. We repeated this process twice per contact map for each simulation configuration (i.e. we sampled twice from the same underlying probability matrix) to generate a pair of “pseudo-replicates”, obtaining 14 (7 x 2) simulated contact maps.

We then simulated various types of noise in the contact maps to evaluate the behavior of the GenomeDISCO concordance score.

- **Edge dropout** The edge dropout simulations measured how our concordance score changes as a function of removing edges from contact map graphs. For this simulation, we randomly set *x*% (for *x* between 10% and 90%) of the entries in the probability matrix to 0. We then renormalized the upper triangle to a valid probability distribution and then sampled from a binomial distribution as described above. For each level of noise, we computed scores by comparing the original sample (0% dropout) against a sample with *x*% edge dropout. We estimated the standard deviation of scores for *x*% edge dropout based on 14 comparisons. Seven of these correspond to concordance scores obtained across the 7 cell types with Hi-C data. Also, for each cell type, we computed 2 scores: one comparison for replicate 1 (0% dropout) vs. replicate 2 (*x*% dropout), the second comparison for replicate 2 (0% dropout) vs. replicate 1 (*x*% dropout).
- **Node dropout** The node dropout simulations involve random removal of nodes. As in the edge dropout simulations, we perturbed *P*. For a given percent of dropout, *x*, we removed *x*% of the nodes, which is equivalent to setting all probabilities involving that node to 0. Then we renormalized and sampled reads from the binomial distribution as described above. The comparisons are analogous to the ones described in the edge dropout simulations: we compared the contact maps with 0% dropout against those with *x*% dropout for a total of 14 comparisons per dropout level.
- **Domain boundary noise** The domain boundary noise simulations were designed to understand how the concordance score changes as a function of uncertainty in the location of domain/TAD boundaries in the contact map. To simulate variation in domain location, we used a reference contact map and shifted it by a specific number of nodes *b* called the domain boundary noise. The contact frequency for a pair of nodes (*i, j*) in the shifted contact map is equal to the contact frequency at nodes (*i* + *b, j* + *b*) in the reference contact map. Then, we compared the original matrix with matrices shifted by different values of *b* (50 kb, 100 kb, 200 kb, 400 kb, 80 kb, 1.6 Mb). We performed this shift using all nodes on chr21, but for scoring, we only used a subset of this chromosome that is contiguous (starting at 20 Mb and ending at 45 Mb), leaving out the rest of the chromosome that is composed of mostly unmappable regions. For consistency with the other simulation types described above, this subset was used in all simulations in this paper. As in the previous simulations, we obtained 14 comparisons per shift.
- **Different distance dependence curves** For each cell type, we rescaled contact maps to simulate each of 7 available distance dependence curves observed in our simulated datasets (Figure S3A), allowing us to test how concordance scores change as a function of differences in distance dependence curves. To transform a contact map to obey a desired distance dependence function, we used the probability matrix *P* representation of the contact map (as described above) and rescaled values at each genomic distance such that the average contact probability at that distance corresponds to the average contact frequency of the desired dataset. Finally, given the rescaled probability matrix, we sampled from the binomial distribution to obtain the simulated datasets. For each of the 7 simulated distance dependence curves, we obtained 2 pseudoreplicates by sampling from the binomial distribution twice. We restricted this experiment to comparisons between samples simulated from the same cell type, where the only variable that changes is the difference in distance dependence curve. To measure the separation between the two groups of comparisons (pairs of samples with the same distance dependence curves and those with different curves), we used a Mann-Whitney test, comparing pairs of contact maps with similar versus different distance dependence curves. Specifically, we split comparisons into two groups based on a threshold difference in distance dependence curve of 0.005 Jensen Shannon divergence (see Section 4), which is approx. half of the largest difference in distance dependence curves we observed. For reference, Figure S3C shows the scores we obtain as a function of difference in distance dependence curves, while Figure 2D in the main text shows the scores grouped by whether they fell in the “similar distance dependence curves” or in the “different distance dependence curves” categories.
- **Comparing simulated pseudoreplicates within a cell type to simulated maps between different cell types** For this simulation, we use the datasets created in the “Edge noise” simulation, with 0% noise. Since each cell type has 2 simulated pseudoreplicates, we can evaluate concordance of pairs of pseudoreplicates from the same cell type and compare it to concordance of pairs of simulated contact maps from different cell types. We measured the difference in score distributions between pseudoreplicates and different cell types using the Mann-Whitney test as described above.

The simulator code is included in the GenomeDISCO package under “simulations_from_real_data.py”.

When running GenomeDISCO on the simulations, we used *t* = 3, as this value was deemed optimal in our parameter optimization.

## 4 Analysis of distance dependence curves and the differences between them

To obtain a quantitative measurement of whether two contact maps have different distance dependence curves, we computed the Jensen Shannon divergence between the two distance dependence curves. Specifically, for each distance dependence curve (composed of the probability of contact as a function of linear genomic distance), we rescale values so that they sum to 1, and then use this probability distribution as input to the Jensen Shannon divergence. These values are computed based on chromosome 21, both for the simulations and for the analysis of real data.

For reference, we include in Figure S3 all distance dependence curves analyzed in the simulations (Figure S3A) and in the real Hi-C data (Figure S3B).

**Figure S3:**
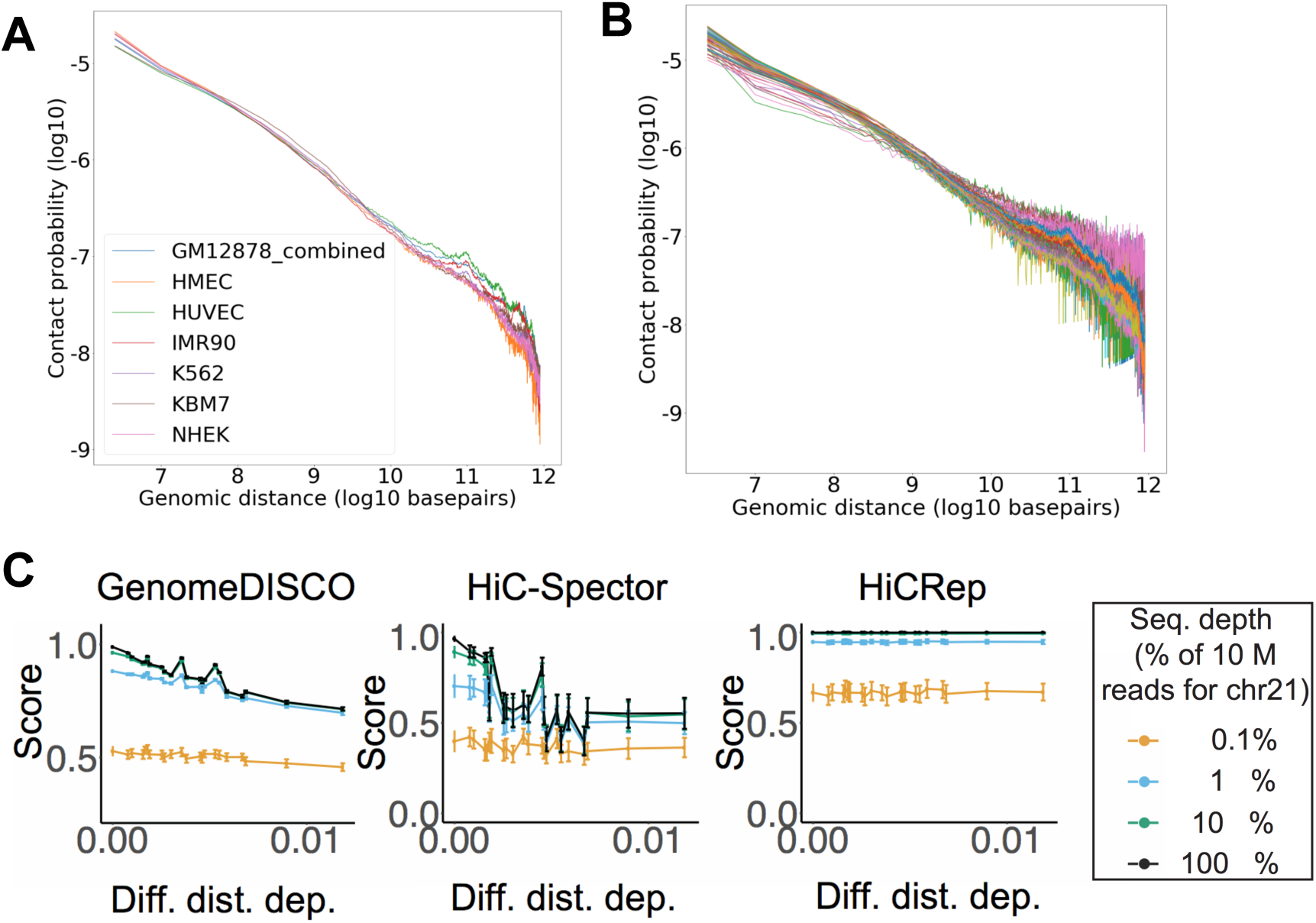
Distance dependence curves. A) All distance dependence curves analyzed in the simulations. B) All distance dependence curves analyzed in the real Hi-C data. C) Scores obtained on the simulations of different distance dependence curves for GenomeDISCO, HiC-Spector and HiCRep. Error bars represent one standard deviation.

## 5 Comparison with other methods

We compared our method with two other recently developed concordance scoring methods for Hi-C data; HiCRep (Yang *et al.*, 2017) and HiC-Spector (Yan *et al.*, 2016). For HiCRep we used a maximum distance of contacts equal to 5 Mb and a smoothing parameter h=5, which is what was suggested for 40kb resolution Hi-C data. For HiC-Spector we used 20 eigenvectors. Another commonly used concordance score is correlation (Spearman or Pearson), but (Yang *et al.*, 2017) have already pointed out its deficiencies. Hence, we do not include comparisons to naive correlation measures.

## 6 Using GenomeDISCO to analyze differences in protocols and restriction enzymes in real Hi-C data

We reasoned that we could use the experiments from (Rao *et al.*, 2014) to investigate how our concordance scores behave when comparing samples prepared with different protocols or different restriction enzymes.

For comparing different protocols (Figure S4A), we filtered out samples that were no-crosslinking controls, or categories with fewer than 3 comparisons. We found that scores comparing in situ Hi-C with dilution Hi-C were lower than for the other protocols, for all 3 methods. This is expected, since (Rao *et al.*, 2014) report that dilution Hi-C contains more contacts between nuclear DNA and mitochondrial DNA, compared to in-situ Hi-C in which the nuclear membrane prevents such random ligations between nuclear and mitochondrial DNA.

**Figure S4:**
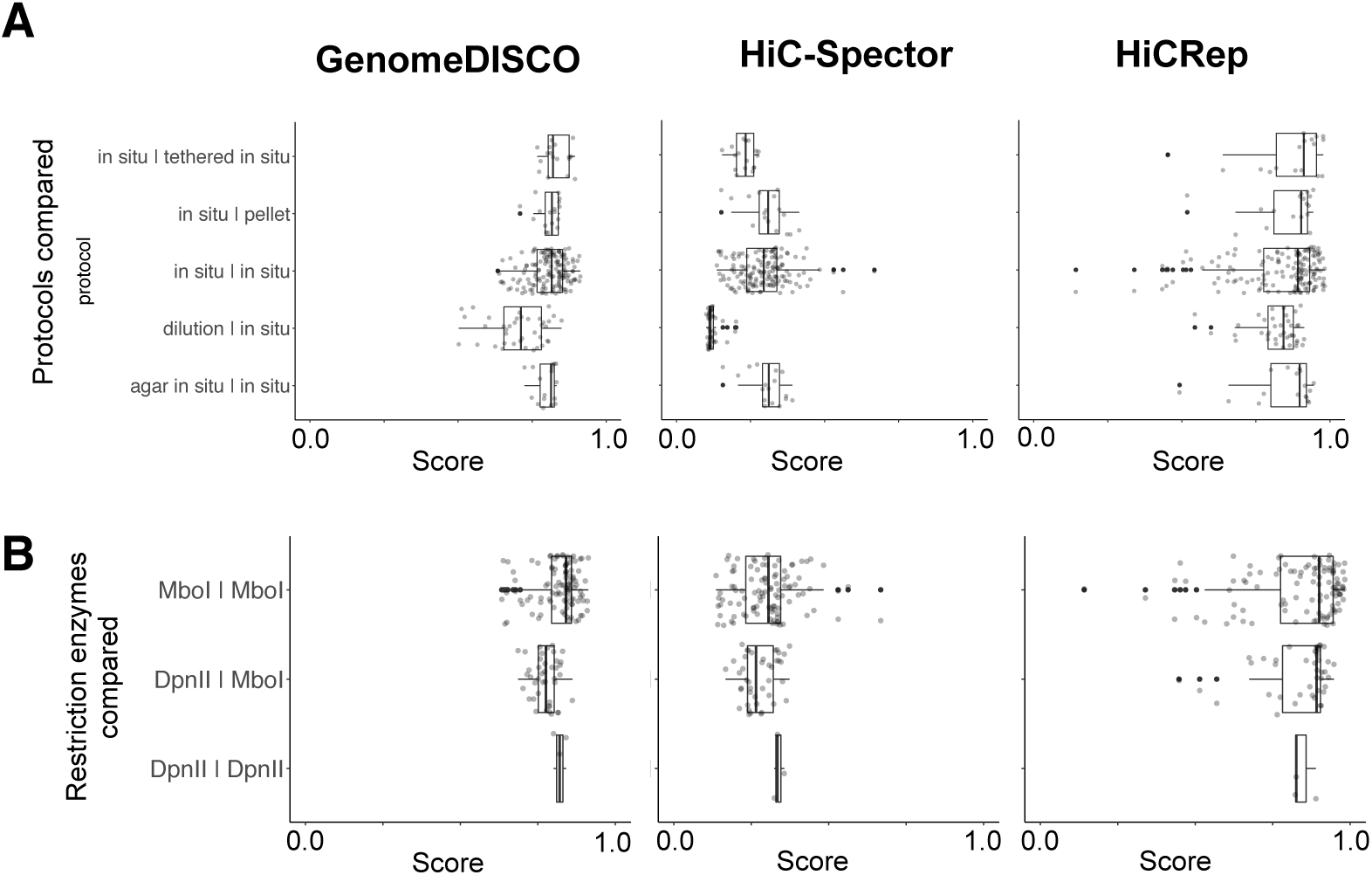
Concordance scores as a function of protocol and restriction enzyme. A) GenomeDISCO, HiC-Spector and HiCRep scores as a function of pairs of protocols compared. B) GenomeDISCO, HiC-Spector and HiCRep scores as a function of pairs of restriction enzymes used.This analysis was restricted to the samples in our test set.

For the comparison of restriction enzymes (Figure S4B), we focused on samples done with in situ Hi-C, to avoid confounding between restriction enzyme used and protocol used. And as before, we required the categories to have a minimum of 3 scores. For the remaining comparisons, GenomeDISCO ranks highest the samples that are prepared with the same restriction enzyme (MboI vs MboI), and assigns lower scores to comparisons done with different restriction enzymes (DpnII vs MboI). The difference between the 2 restriction enzyme combinations is as expected small, since both DpnII and MboI are 4-cutters.

## 7 Using GenomeDISCO to cluster experiments

For the analysis of real Hi-C data from (Rao *et al.*, 2014), we asked whether we could cluster the samples based on the matrix of scores obtained from all pairwise comparisons between the samples in our test set. For this, we started from a matrix of concordance scores and performed hierarchical clustering on this matrix, with a Euclidean distance measure and complete linkage. The resulting dendrograms are in Supplementary Figure 5A. As expected, all methods in general produce clusters of biological replicates. To quantify this, we considered each cut in the hierarchical clustering (in other words, each number of clusters, from 1 to the number of samples), and for each cut computed an adjusted Rand index to quantify whether the clustering at that stage partitions samples from the same cell type in the same cluster (Supplementary Figure 5B). We find that GenomeDISCO achieves the highest maximum adjusted Rand index. Interestingly, both GenomeDISCO and HiCRep produce dendrograms in which one sample from the GM12878 cell type (sample HIC048) does not cluster with the rest of the samples from the same cell type. This is consistent with the observation from (Rao *et al.*, 2014) that sample HIC048 is from a different batch of cells than the rest of the GM12878 samples in the dendrogram. Specifically, it is from “Batch 1, received at the Broad Institute on 3/11/2008”, whereas the other samples are from “Batch 2, received at the Broad Institute on 5/13/2008”. Cells from batch 2 were found to harbor a translocation between chromosomes 6 and 11, which was not present in batch 1 in (Rao *et al.*, 2014). Thus, GenomeDISCO was able to correctly identify this batch effect related to structural variation.

**Figure S5:**
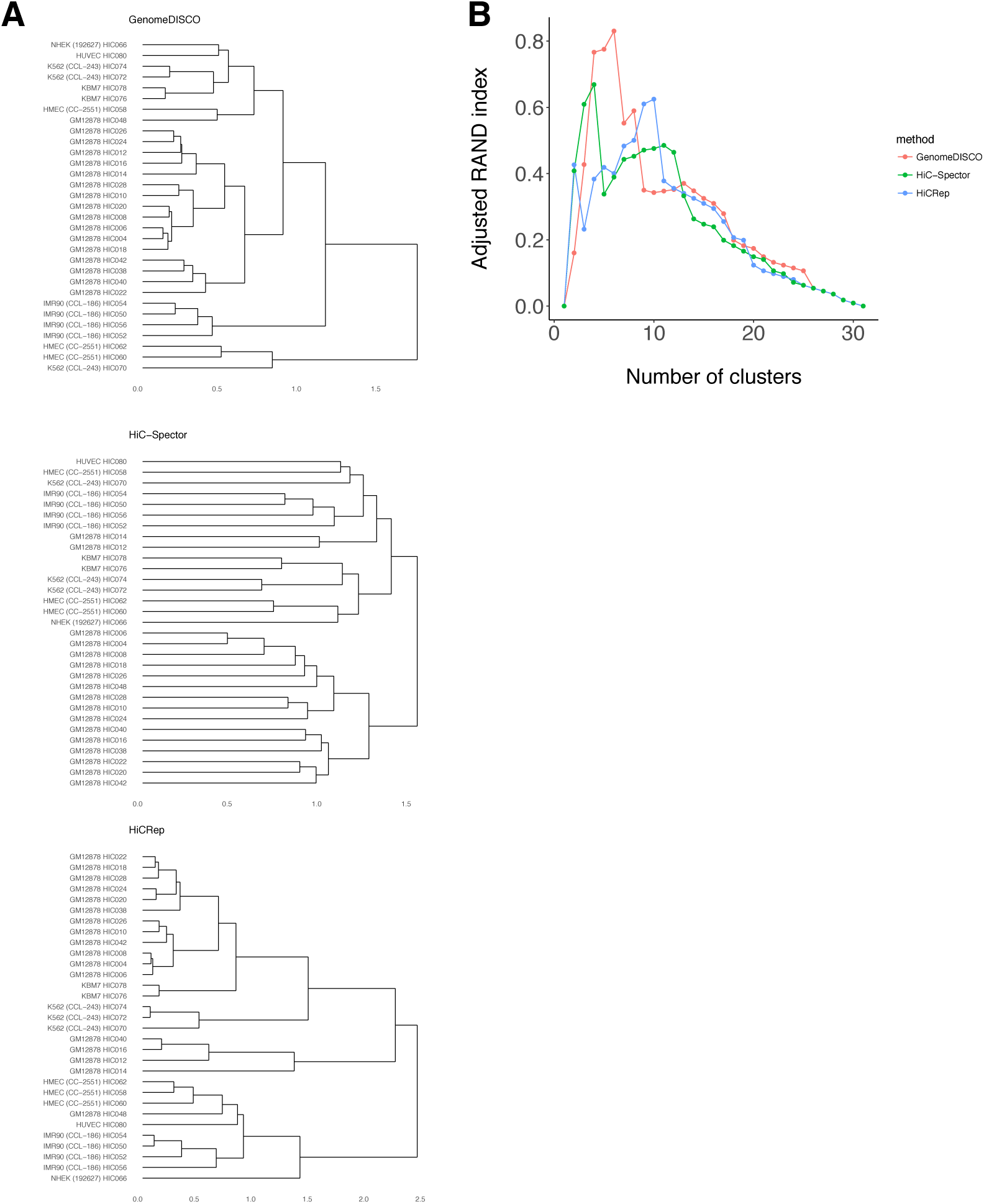
Clustering samples based on their concordance score profiles. A) Dendrograms obtained from clustering samples from (Rao *et al.*, 2014) for GenomeDISCO, HiCRep and HiC-Spector, based on the similarity matrix obtained from all pairwise concordance scores. B) Adjusted RAND scores for each possible number of clusters (obtained from cutting the hierarchical clustering tree at each possible level). This analysis was restricted to the samples in our test set.

## 8 Running time analysis

We profiled the running time and memory requirements for GenomeDISCO and the other methods compared here. We ran each method genomewide (i.e. sequentially across all chromosomes) on 50 different comparisons from Rao *et al.* (2014), so that we could estimate the variance of the running times. All analyses were run on an Intel Xeon CPU E5-2683 v3 running at 2.00GHz, and we report as running times the wall clock times measured with the “time” command in Unix.

We found that HiC-Spector was the fastest (median running time 5.02 minutes), followed by HiCRep (median running time 17.87 minutes) and GenomeDISCO (median running time 31.49 minutes), as we show in Figure S6. GenomeDISCO displayed the largest variance in running time because the main operation involves the multiplication of contact maps during the random walk calculation, and given that the contact maps are stored as sparse matrices, the running time depends on the level of sparsity of the matrices. On the other hand, HiCRep and HiC-Spector operate on fixed size inputs, with HiC-Spector focusing on the top 20 eigenvectors, and HiCRep working with the 2D matrix which has a fixed size. Because GenomeDISCO’s running time depends on the sparsity of the matrices, it was observed to run faster than HiCRep in Yardimci *et al.* (2017), where the datasets used for benchmarking running times are sequenced at less than 50 million reads, compared to the datasets considered in this paper that are sequenced more deeply.

**Figure S6:**
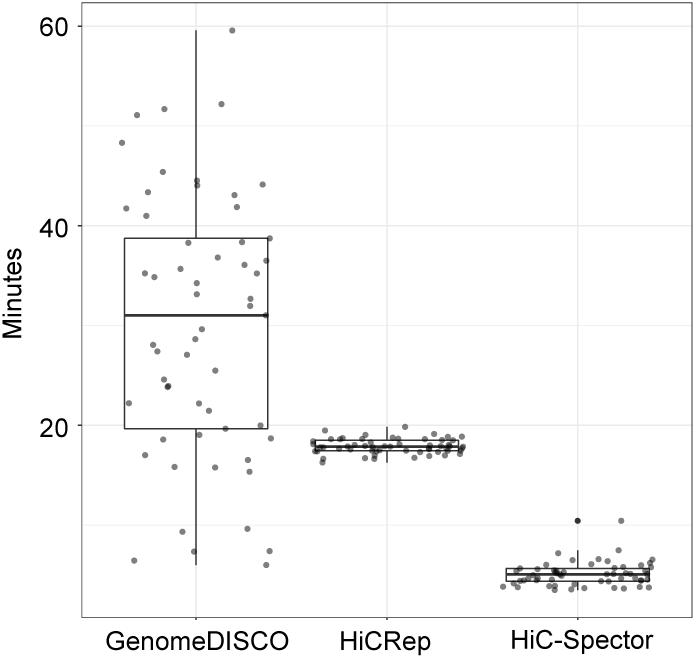
Running times of GenomeDISCO, HiCRep and HiC-Spector.

